# Using large-scale mutagenesis to guide single amino acid scanning experiments

**DOI:** 10.1101/140707

**Authors:** Vanessa E. Gray, Ronald J. Hause, Douglas M. Fowler

## Abstract

Alanine scanning mutagenesis is a widely-used method for identifying protein positions that are important for function or ligand binding. Alanine was chosen because it is physicochemically innocuous and constitutes a deletion of the side chain at the β- carbon. Alanine is also thought to best represent the effects of other mutations; however, this assumption has not been formally tested. To determine whether alanine substitutions are always the best choice, we analyzed 34,373 mutations in fourteen proteins whose effects were measured using large-scale mutagenesis approaches. We found that several substitutions, including histidine and asparagine, are better at recapitulating the effects of other substitutions. Histidine and asparagine also correlated best with the effects of other substitutions in different structural contexts. Furthermore, we found that alanine is among the worst substitutions for detecting ligand interface positions, despite its frequent use for this purpose. Our work highlights the utility of large-scale mutagenesis data and can help to guide future single substitution mutational scans.

## Introduction

Making and studying mutants is a fundamental way to learn about proteins, revealing functionally important positions, validating specific hypotheses about catalytic mechanism and yielding insights into protein folding and stability. Single amino acid scanning mutagenesis, in which every position in a protein is sequentially mutated to one particular amino acid, was a key advance. By searching sequence space systematically, scanning mutagenesis enabled the unbiased identification of positions important for protein function. The first application of scanning mutagenesis used alanine substitutions to identify positions in human growth hormone important for receptor binding^1^. Alanine was chosen because it represents a deletion of the side chain at the β-carbon, and because, being uncharged and of modest size, it is physicochemically innocuous. Furthermore, alanine is the second most abundant amino acid in natural sequences and is found in a variety of structural contexts^2–4^. In addition to alanine, many other amino acids including arginine^5^, cysteine^6^, glycine^7^, methionine^8^, phenylalanine^9^, proline^10^ and tryptophan^11^ have been used for scanning mutagenesis, often with a specific hypothesis in mind *(e.g*. that bulky amino acids are important). Nevertheless, the vast majority of scanning mutagenesis experiments are conducted using alanine under the assumption that alanine substitutions are especially useful for identifying functionally important positions.

Does alanine best represent the effect of other substitutions? Are alanine substitutions ideal for finding functionally important positions, such as those that participate in binding interfaces? Answering these questions is important because alanine scans continue to be used to understand and engineer proteins. Despite the large investment in alanine scanning mutagenesis, little work has been done to determine which substitutions are ideal. Some scanning mutagenesis studies compare two different types of scans *(e.g*. alanine and cysteine), but generally find that the information revealed by each substitution is distinct^9,12^. Computational predictions for all substitutions at 1,073 positions across 48 proteins in the Alanine Scanning Energetics Database suggested that alanine substitutions correlated best with the mean effect of every mutation at each position^13^. However, concrete answers to these questions require comparing the empirical effects of different substitutions in many proteins. Thus, we analyzed large-scale experimental mutagenesis data sets comprising 34,373 mutations in fourteen proteins. We found that proline is the most disruptive substitution and methionine is the most tolerated. Global and position-centric analyses revealed that histidine and asparagine substitutions best represent the effects of other substitutions. We evaluated the utility of each amino acid substitution for determining whether a position is in a ligand-binding interface, and found that aspartic acid, glutamic acid, asparagine and glutamine performed best. Thus, our results suggest that, compared to other substitutions, alanine substitutions are not especially representative, nor are they the best choice for finding ligand-binding interfaces.

## Results

Deep mutational scanning is a method that enables measurement of the effects of hundreds of thousands of mutations in a protein simultaneously^14,15^. Deep mutational scanning can be used to quantify the effects of all mutations at each position in a protein, and is therefore a conceptual extension of single amino acid scanning mutagenesis. Broad application of deep mutational scanning has resulted in an explosion of protein mutagenesis data^14^. These large-scale mutagenesis data sets create the opportunity to assess relationships between the effects of different amino acid substitutions comprehensively.

We curated sixteen large-scale mutagenesis data sets from published deep mutational scans of fourteen proteins (Fig. 1A, Table 1). Here, we included two distinct data sets for the BRCA1 RING domain and for UBI4 because mutations in these proteins have been independently assayed for different protein functions (*e.g*. BRCA1 BARD1 binding and E3 ligase activity). Our collection of data sets is ideal for an unbiased analysis of the general effects of mutations because the mutagenized proteins are highly diverse, encompassing enzymes, structural proteins and chaperones from organisms ranging from bacteria to humans. The frequency of amino acids in the wild type sequences of the fourteen proteins was similar to amino acid frequencies in all known proteins^2^ (Fig. 1B). For example, leucine (frequency = 11%) and alanine (8%) were the most frequently occurring wild type amino acids in the fourteen proteins, while tryptophan (<1%) was the rarest. However, the unbiased and massively parallel nature of deep mutational scanning experiments yielded a relatively uniform distribution of amino acid substitutions (Fig. 1C). Furthermore, the data sets were generated by different labs at different times using different types of assays, reducing the chances of bias arising from specific experimental or analytical practices. Importantly, the assay formats used for the deep mutational scans included many commonly employed in alanine scanning like phage display and yeast two-hybrid. Collectively, these large-scale mutagenesis data sets comprised 34,373 nonsynonymous mutations at 2,236 positions in the fourteen proteins. The data sets contained effect scores for most mutations at each position. To facilitate comparisons between each data set, we rescaled mutational effect scores for each protein, using synonymous mutations to define wild type-like activity and the bottom 1% of mutations to define lack of activity (Fig. S1A). Thus, each mutational effect score reflects the impact of the mutation, relative to wild type, with a score of zero meaning no activity and a score of one meaning wild type-like activity.

**Figure 1.**
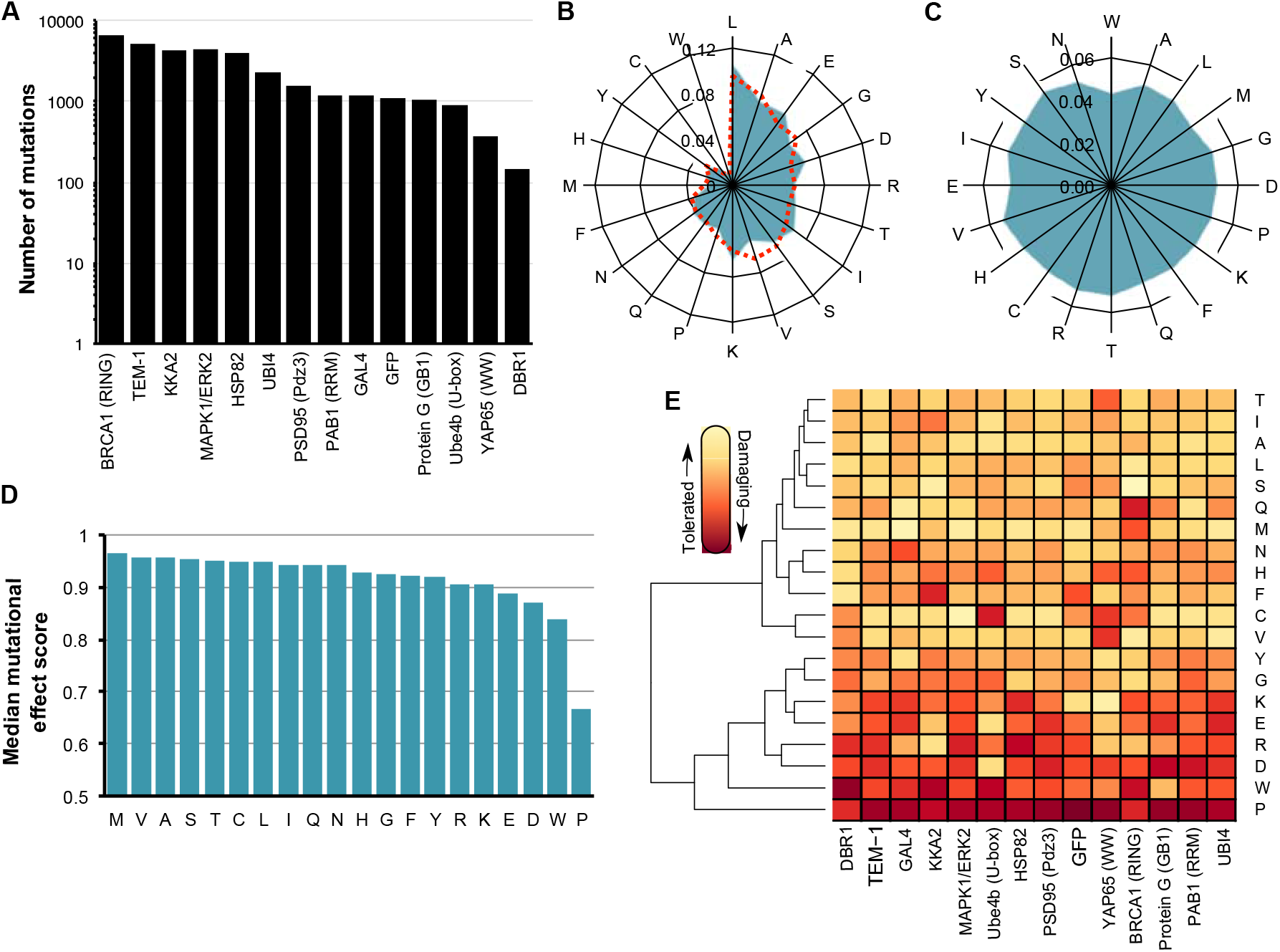
Large-scale mutagenesis data from fourteen proteins. **(A)** The number of single amino acid mutations with effect scores in each of the fourteen proteins is shown. **(B)** A radar plot shows the relative frequency of occurrence of each amino acid in the wild type sequences of the fourteen proteins (blue) or in 554,515 proteins in the UniProt Knowledgebase^2^ (dashed red). **(C)** A radar plot shows the relative frequency of each of the twenty amino acid substitutions in the large-scale mutagenesis data sets for all fourteen proteins. **(D)** The median mutational effect score of each amino acid substitution is shown for 34,373 mutations at 2,236 positions in all fourteen proteins. **(E)** A heat map shows the median mutational effect score of each amino acid substitution for each protein separately. Yellow indicates tolerated substitutions while orange indicated damaging substitutions. Amino acids and proteins were ordered according to similarity using hierarchical clustering with the hclust function from the heatmap2 package in R. The dendrogram is shown only for amino acid clustering.

**Table 1.** Large-scale mutagenesis data sets used in this study

| Data set | Number of mutations | Mutagenized positions | Mutational completeness* | Organism | Selected Phenotype | Citation |
| --- | --- | --- | --- | --- | --- | --- |
| Aminoglycoside kinase | 4,234 | 264 | 84% | *K. pneumoniae* | Antibiotic resistance | 31 |
| BRCA1 RING domain - BARD1 binding | 1,748 | 102 | 90% | *H. sapiens* | Binding activity (Y2H) | 28 |
| BRCA1 RING domain - E3 ligase activity | 4,872 | 303 | 85% | *H. sapiens* | Ubiquitin ligase activity | 28 |
| DBR1 | 144 | 25 | 30% | *H. sapiens* | RNA enzyme activity | 32 |
| Gal4 | 1,196 | 64 | 98% | *H. sapiens* | Transcription factor activity | 33 |
| GFP | 1,084 | 235 | 24% | *A. victoria* | Fluorescence | 34 |
| Hsp82 | 4,021 | 219 | 97% | *S. cerevisiae* | Chaperone activity | 35 |
| hYAP65 WW domain | 363 | 33 | 58% | *H. sapiens* | Ligand binding | 15 |
| MAPK1/ERK2 | 4,470 | 359 | 66% | *H. sapiens* | Kinase activity | 36 |
| Pab1 | 1,188 | 75 | 83% | *S. cerevisiae* | mRNA binding | 37 |
| Protein G GB1 domain | 1,045 | 55 | 100% | *Streptococcus sp. group G* | IgG-Fc binding | 25 |
| PSD95 pdz3 domain | 1,577 | 83 | 100% | *H. sapiens* | Ligand binding | 26 |
| TEM1 β-lactamase | 5,198 | 287 | 95% | *E. coli* | Antibiotic resistance | 38 |
| Ube4b U-box | 899 | 102 | 46% | *S. cerevisiae* | Ubiquitin ligase activity | 39 |
| Ubi4 - activity | 1,249 | 75 | 88% | *S. cerevisiae* | Ubiquitin ligase activity | 40 |
| Ubi4 – activation by E1 | 1,085 | 60 | 95% | *S. cerevisiae* | Ubiquitin ligase activity | 41 |

\*proportion of all possible single amino acid mutations in mutagenized region observed

To validate the large-scale mutagenesis data, we examined expected patterns of mutational effect. For example, mutations to proline should generally disrupt protein function, as proline restricts the conformation of the polypeptide backbone and eliminates the amide hydrogen necessary for hydrogen bonding. Indeed, proline substitutions were overwhelmingly more damaging than other substitutions to protein function (Fig. 1D; Fig.S1B). In fact, proline was the most damaging amino acid in eleven of fourteen proteins and second most damaging in the remaining three proteins (Fig. 1E). Additionally, as expected from the Dayhoff^16^, Blosum^17^ and Grantham^18^ substitution matrices, tryptophan tended to be deleterious. Methionine was the best-tolerated substitution. Many other substitutions were also well-tolerated, with seven different amino acids appearing as the most tolerated across the fourteen proteins (Fig. 1D, E). Tolerance to substitutions depends on structural context, so the variability in the best tolerated substitution might be due to diversity in the structural composition of each protein in our data set. Thus, the large-scale mutagenesis data sets we collected generally recapitulated our expectations about the effects of mutations, despite coming from fourteen distinct proteins that were each assayed independently.

Next, we determined which amino acid substitution best represented the effects of all other substitutions. To avoid bias arising from incomplete data, we restricted this analysis to the 882 positions in the fourteen proteins with measured effects for all nineteen possible substitutions. We calculated the median mutational effect at each of these 882 positions. Overall, the median effects across these positions were mildly damaging, with a mean of 0.82 (stop ~ 0, wild-type ~ 1). We found that the effects of phenylalanine, glycine, histidine, isoleucine, leucine, asparagine, glutamine and tyrosine substitutions were all indistinguishable from the median effects (Fig. 2A, **Table S1**). However, proline, aspartic acid and tryptophan substitutions are much more damaging than the median substitution. Alanine, cysteine, methionine, serine, threonine and valine are considerably less damaging than the median substitution. These well-tolerated amino acid substitutions might be useful for detecting the most mutationally sensitive positions in a protein. However, these substitutions, alanine included, are not especially representative of the effects of other substitutions.

**Figure 2.**
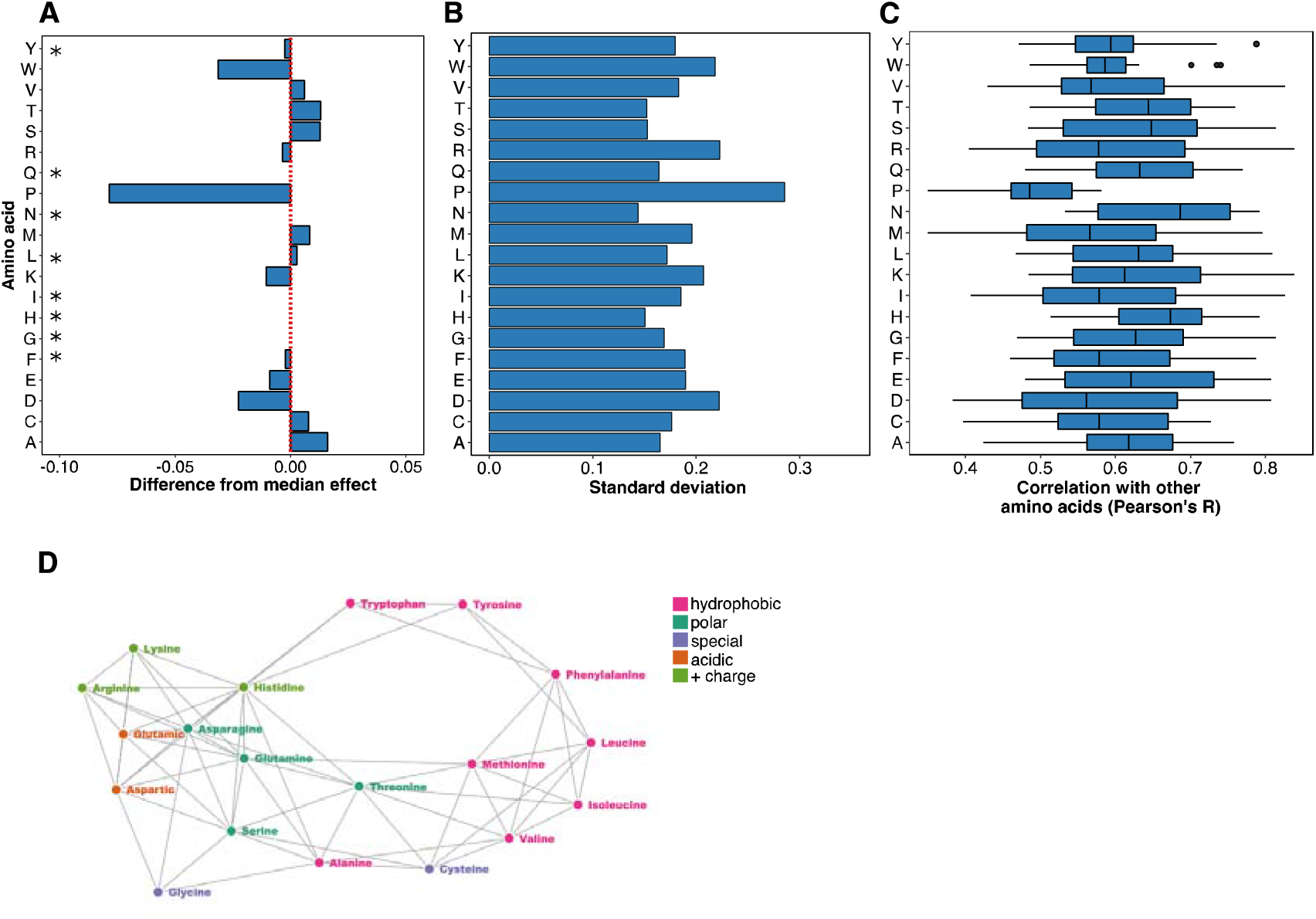
Histidine and asparagine substitutions best represent the effect of other substitutions. **(A)** For each of the 882 positions where the mutational effects of all nineteen substitutions were measured, the difference from the median effect was calculated for each substitution at each position. The median of these differences across all positions for each substitution is shown, with the red line indicating a median difference of zero. A paired, two-sided Wilcoxon rank sum test was used to determine whether each substitution’s difference from the median effect across all positions was equal to zero (* indicates substitutions with a Bonferroni-corrected *p*-value > 0.01; Table S1). **(B)** The standard deviation of each substitution’s differences from the median effect at the 882 positions where the mutational effects of all nineteen substitutions were measured is shown. **(C)** For each substitution, Pearson correlation coefficients were calculated for the mutational effects of that substitution with every other substitution at each position. The distribution of correlation coefficients for each substitution is shown. **(D)** These pairwise mutational effect score correlations are also illustrated using a force directed graph. Each node represents an amino acid and each edge force value is the Pearson correlation coefficient for the mutational effect scores of the two amino acid substitutions connected by the edge. To reduce the density of edges, only the top 40% of Pearson correlation coefficients were included. This cutoff removed proline from the graph. Amino acids are colored by physicochemical type. The graph was constructed using the networkD3 package in R.

We also examined the dispersion of each amino acid’s mutational effect about the median at all 882 positions, reasoning that representative substitutions would have minimal dispersion. Of substitutions whose effects were indistinguishable from the median effect, histidine and asparagine have the smallest dispersion (standard deviation = 0.15 and 0.14, respectively; Fig. 2B), while tyrosine (0.18), glutamine (0.16), phenylalanine (0.19), glycine (0.17), leucine (0.17) and isoleucine (0.19) all had larger dispersions. Thus, of all possible substitutions, histidine and asparagine tended to have effects closest to the median effect at the 882 positions we examined.

Because of the comprehensive nature of the large-scale mutagenesis data sets, we could ask how well the mutational effect scores of each substitution correlated with the scores of every other substitution at each position. Thus, we calculated Pearson correlation coefficients for the mutational effect scores of each substitution pair across all positions (Fig. 2C, Fig. S2). The effects of histidine and asparagine substitutions correlated best with the effects of all other substitutions, while the effect of proline substitutions correlated worst. To visualize the relationships between each pair of substitutions, we constructed a force-directed graph (Fig. 2D). As expected, substitutions cluster by physicochemical type in the graph, meaning that similar substitutions have similar effects. Proline is not represented because its effects are poorly correlated with other substitutions. Histidine and asparagine are connected to many other amino acids, owing to the high correlation of the effects of these substitutions with many other substitutions.

We next asked whether the secondary structural context of a position altered the effect of each substitution. We excluded DBR1 and GB1 from this analysis because they did not have structures of a sufficiently close homologs. We used DSSP to identify 1,007 positions in the remaining proteins that were in an α-helix, a β-sheet or a turn^19^. Overall, substitutions in turns are less damaging than substitutions in α-helices or β-sheets (Fig. 3A). However, the relative effects of each substitution in the three structural contexts were mostly consistent, especially between α-helices and β-sheets (Fig. 3B, S3A). Surprisingly, the tolerance for each amino acid substitution in the different secondary structural contexts was not strongly correlated with the frequency of that amino acid’s occurrence in known structures^20^. For example, alanine occurs more frequently in α- helices, relative to β-sheets. However, in our large-scale mutagenesis data sets, alanine substitutions were mildly damaging in both structural contexts. These observations suggest that secondary structure does not dominate mutational tolerance, at least for the proteins we examined.

**Figure 3.**
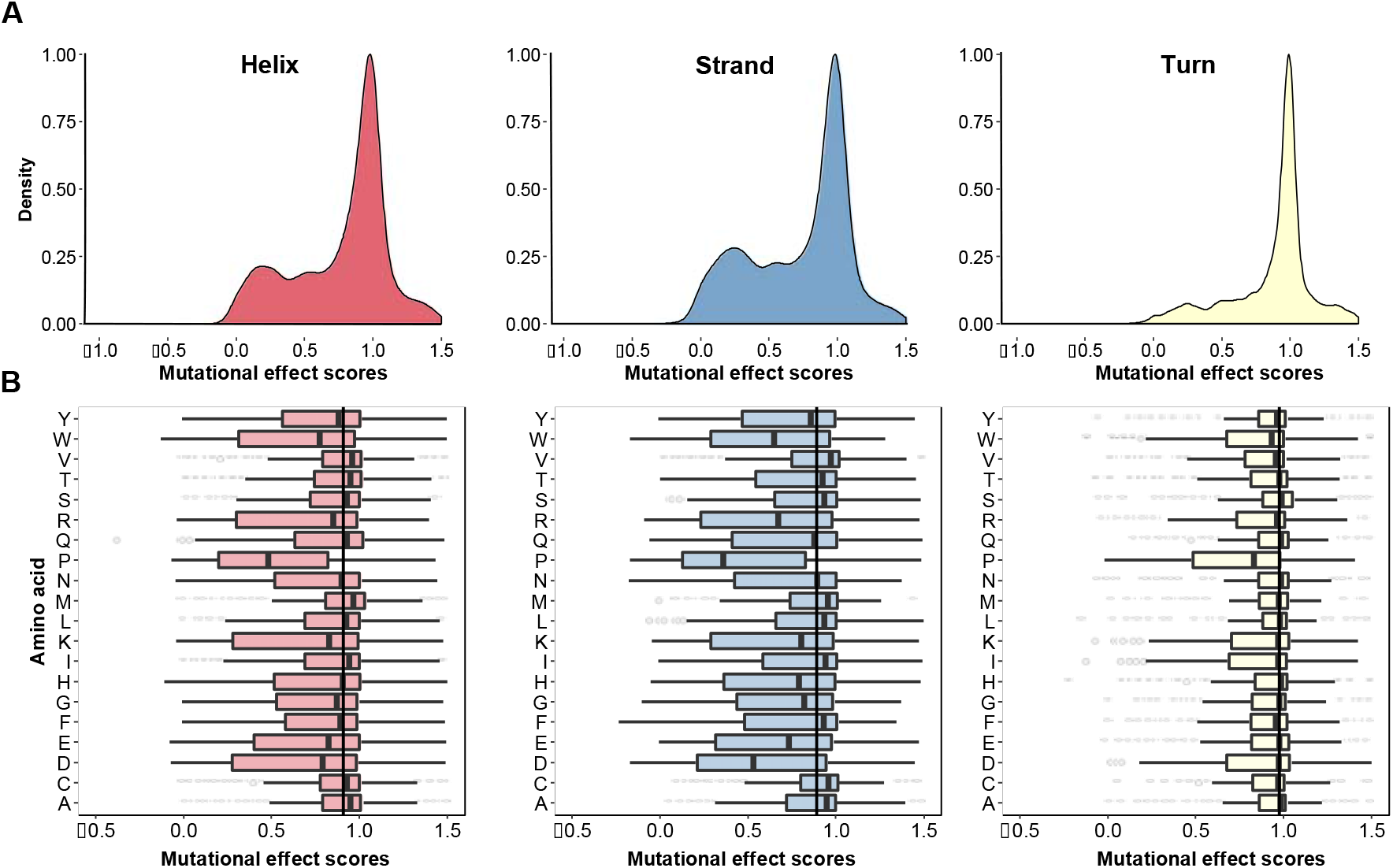
Secondary structural context of mutational effects. **(A)** Density functions describing the distribution of mutational effect scores for each substitution are shown for three different structural contexts as determined using DSSP: β-sheets (left panel, N = 4,796), α-helices (middle panel, N = 8,669) and turns (right panel, N = 3,329). **(B)** The mutational effect score distributions for each substitution in β-sheets (left panel), α- helices (middle panel), and turns (right panel) are shown. The vertical line in each panel represents the median effect score for all substitutions in that secondary structure type.

We next investigated which substitutions were the most representative regardless of structural context. We found that histidine substitutions have close to the median effect in α-helices and turns, but were more damaging than the median effect in β-sheets (Fig. 3B). Asparagine and glutamine substitutions had near median effects in all three contexts. As above, we examined how well the effects of each substitution correlated with every other substitution at each position in each context. We found that the effects of histidine, asparagine and glutamine substitutions correlated best with the effects of other substitutions (Fig. S3B, C). Thus, the effects of histidine, asparagine and glutamine are relatively consistent in the different structural contexts we examined, highlighting the representativeness of these substitutions.

An important use of alanine scanning is to identify positions in protein-ligand interfaces. In order to determine whether alanine is ideal for that purpose, we analyzed the effects of substitutions in four proteins with ligand-bound structures: the hYAP65 WW domain, the PSD95 pdz3 domain, the BRCA1 RING domain and GAL4. Amongst these four proteins there were 4,884 mutations at 282 positions. We used relative solvent exposure to classify each position as either buried or on the surface. We also determined interface positions based on published structures and functional studies (see Methods). We found that substitutions at interface positions are substantially more damaging than substitutions at buried, non-interface or surface, non-interface positions (Fig. 4A). This result is expected, given that all four deep mutational scans were conducted using selections that depended on ligand binding. Alanine, along with phenylalanine, isoleucine and methionine, are the least disruptive amino acid substitutions at interface positions, suggesting that they may not be ideal for interface detection.

**Figure 4.**
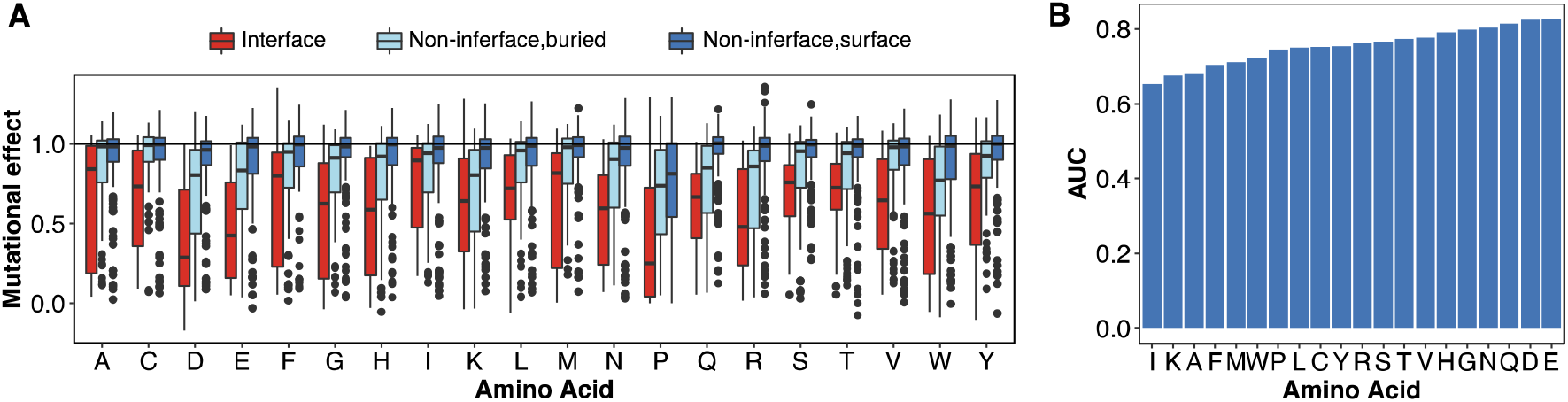
Alanine is not especially useful for identifying positions in protein-ligand interfaces. **(A)** The distribution of mutational effect scores for every substitution in four proteins with ligand-bound structures (hYAP65 WW domain, PSD95 pdz3 domain, BRCA1 RING domain (BARD1 binding) and Gal4) is shown at ligand interface positions as reported in the literature, and for non-interface buried positions or non-interface surface positions. **(B)** A mutational effect threshold was defined such that positions with a mutational effect below the threshold were classified as “interface,” whereas positions with a mutational effect above the threshold were classified as “non-interface.” ROC curves for each amino acid were generated by varying this threshold. The area under each ROC curve is shown, illustrating the power of each substitution to discriminate between interface and non-interface positions.

We reasoned that the ideal substitution for detecting protein-ligand interfaces would exhibit a large difference in mutational effect between interface and non-interface positions. To formalize this idea, we used a mutational effect threshold. If a substitution at a particular position had a mutational effect below the threshold, we classified that position as “interface.” Conversely, if the mutational effect was above the threshold that position was classified as “non-interface.” For each substitution, we varied the mutational effect threshold from the maximum mutational effect score to the minimum effect in 200 steps. At each step, we compared the true interface positions to those determined using the mutational effect threshold procedure. We then constructed receiver operating characteristic (ROC) curves. The area under each ROC curve revealed the ability of that substitution to discriminate between true interface and non-interface positions. Surprisingly, we found that alanine had among the worst discriminatory power (Fig. 4B, Fig. S4). Substitutions that were highly disruptive at interfaces, like asparagine, glutamine, aspartic acid and glutamic acid, had the best discriminatory power. Next, we calculated the fraction of true interface positions detected by each amino acid substitution at a 5% false positive rate. Here, we found that asparagine and glutamine substitutions revealed over 60% of the true interface positions; aspartic acid and glutamic acid substitutions also performed well (Fig. S5). However, alanine substitutions detected fewer than 20% of the true interface positions at a 5% false positive rate. Thus, asparagine, glutamine, aspartic acid or glutamic acid substitutions are all better choices than alanine for detecting protein-ligand interfaces.

## Discussion

Alanine scanning mutagenesis is a widely-used method for identifying protein positions that are important for function or ligand binding. Alanine was selected on rational grounds: it is physicochemically innocuous and constitutes a deletion of the side chain at the β-carbon. By analyzing tens of thousands of mutations in fourteen proteins, we have determined that alanine is not the most revealing substitution. In fact, many superior choices exist. For example, histidine and asparagine substitutions have an effect close to the median, and these substitutions correlate best with the effects of all other substitutions. Thus, they better represent the effects of mutations generally.

Asparagine, glutamine, aspartic acid and glutamic acid are the most useful substitutions for detecting ligand interface positions. Thus, our work highlights the utility of large-scale mutagenesis data and suggests that alanine is not necessarily the best choice for future single substitution mutational scans whose goal is to identify functionally important positions or map protein-ligand interfaces.

However, our conclusions are based on only fourteen proteins. While these proteins are diverse in structure and function, they may not fully reflect the mutational propensities of other proteins. For example, tryptophan scanning mutagenesis is often applied to transmembrane domains^21–23^, which were absent from the proteins we analyzed. Thus, our conclusions are most applicable to soluble proteins. Furthermore, we do not address specialized applications of single amino acid scanning mutagenesis. For example, cysteine scanning mutagenesis has been used to introduce disulfide bridges^6^ and glycine scanning mutagenesis has been used to increase conformational flexibility^24^. Our conclusions do not apply to these situations. Finally, the deep mutational scanning data we analyzed arises from genetic selections for protein function. Biochemical assays might reveal different patterns. However, we note that a few of the large-scale mutagenesis data sets we used were benchmarked against and found to be consistent with biochemical assay results^25,26^.

Deep mutational scanning can reveal the functional consequences of all possible single amino acid substitutions in a protein. However, these experiments can be expensive or unwieldy. Therefore, scanning mutagenesis with one or a few amino acids will remain useful for determining functionally important positions, probing protein-ligand interactions and answering other specific questions. Our results could be used to guide future single amino acid scanning mutagenesis experiments, enabling selection of the amino acid best suited for the goals of the experiment.

## Acknowledgements

This work was supported by the National Institute of General Medical Sciences (1R01GM109110 to D.M.F.). V.E.G. is a National Science Foundation Graduate Research Fellow (DGE-1256082) and R.J.H. is a Damon Runyon Cancer Research Foundation Fellow (DRG-2224-15). We thank Lea Starita for helpful discussions and advice.

## Author Contributions

D.M.F. conceived of the project. V.E.G. and R.J.H. curated and rescaled the data sets. V.E.G. and D.M.F. analyzed the data and wrote the paper.

## Materials and Methods

### Data curation and rescaling

We curated a subset of the published deep mutational scanning data sets. We excluded deep mutational scans of non-natural proteins, because the mutational properties of natural and non-natural proteins could differ. The result was a set of sixteen deep mutational scans of fourteen proteins (Table 1). BRCA1 and UBI4 each have two large-scale mutagenesis data sets corresponding independent experiments in which different functions were assayed *(e.g*. ligand binding or catalytic activity). We treated these data sets separately, and did not perform any averaging of mutational effects between the data sets. Additionally, we removed any variants with more than one amino acid substitution from all the data sets.

Most of the data sets reported mutational effect scores as the log-transformed ratio of mutant frequency before and after selection, divided by wild type frequency before and after selection. For data sets that used a different scoring scheme, we recalculated mutational effect scores as the log-transformed ratio of mutant frequency before and after selection, divided by wild type frequency before and after selection. Given that the assays used to detect mutational effect differ, we rescaled the reported mutational effect scores for each data set. First, we subtracted the median effect of synonymous mutations from each reported effect score and then divided by the negative of the bottom 1% of reported effect scores. Finally, we added 1. In cases where synonymous mutational effect scores were unavailable, we omitted the synonymous score median subtraction step. Our rescaling scheme is expressed as

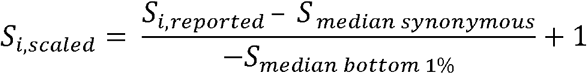

where S is the mutational effect score. Our normalization scheme resulted in scaled mutational effect scores where the most damaging mutations have effect scores ≈ 0 and wild-type-like mutations have scores ≈ 1.

Unless otherwise stated, we used all of the rescaled mutational effect data for each analysis. In each analysis, we used median as a summary statistic rather than mean because the frequency distributions of mutational effect are bimodal rather than Gaussian (Fig. S1).

### Variant annotation

DSSP was used to annotate the secondary structure and absolute solvent accessibility of each wild type amino acid in our data set (http://swift.cmbi.ru.nI/gv/dssp/DSSP_3.html). To estimate the relative solvent accessibility of amino acids, we divided absolute solvent accessibility as determined using DSSP by the total surface area of each amino acid. Amino acids with relative solvent accessibilities greater than 0.2 were labeled as “surface”, whereas amino acids with relative solvent accessibilities less than 0.2 were labeled as “buried”^27^.

### Identification of interface positions

Four proteins in our data set had high-resolution PDB structures with peptide or nucleotide ligands, Gal4 (3COQ), BRCA1 RING domain (1JM7), PSD95 pdz3 domain (1BE9) and hYAP65 WW domain (1JMQ). We determined interface positions from the literature^15,28–30^. The interface positions in hYAP6_5_ WW domain were 188, 190, 197 and 199. The interface positions in BRCA_1_ RING domain were 11, 14, 18, 93 and 96. PSD95 pdz3 domain positions were 318, 322–327, 329, 339, 372 and 379. Gal4 interface positions were 9, 15, 17, 18, 20, 21, 43, 46 and 51.

### Construction of ROC curves

We constructed empirical ROC curves to illustrate the power of each substitution to discriminate between interface and non-interface positions, determined as described above. First, we defined a discrimination threshold, such that positions with a mutational effect score below the threshold were classified “interface” and positions with a mutational effect score above the threshold were classified as “non-interface.” For each substitution, we varied this discrimination threshold from the maximum mutational effect score to the minimum mutational effect score in 200 steps, calculating the true positive interface detection rate (TPR) and false positive interface detection rate (FPR) at each step. The TPR was calculated by dividing the number of interface positions with scores below the mutational effect threshold by the total number of interface positions. The FPR was calculated by dividing the number of non-interface positions with scores below the mutational effect threshold by the total number of non-interface positions. ROC curves were constructed by plotting the TPR and FPR for each of the 200 mutational effect thresholds. The area under each ROC curve was determined in R using the auc() function in the pROC package (https://cran.r-project.org/web/packages/pROC/pROC.pdf)

### Data and code availability

The data sets used in this study came from a variety of published works (see Table 1). The curated data sets and code for generating figures can be found at: https://github.com/FowlerLab/

---

**Figure S1.**
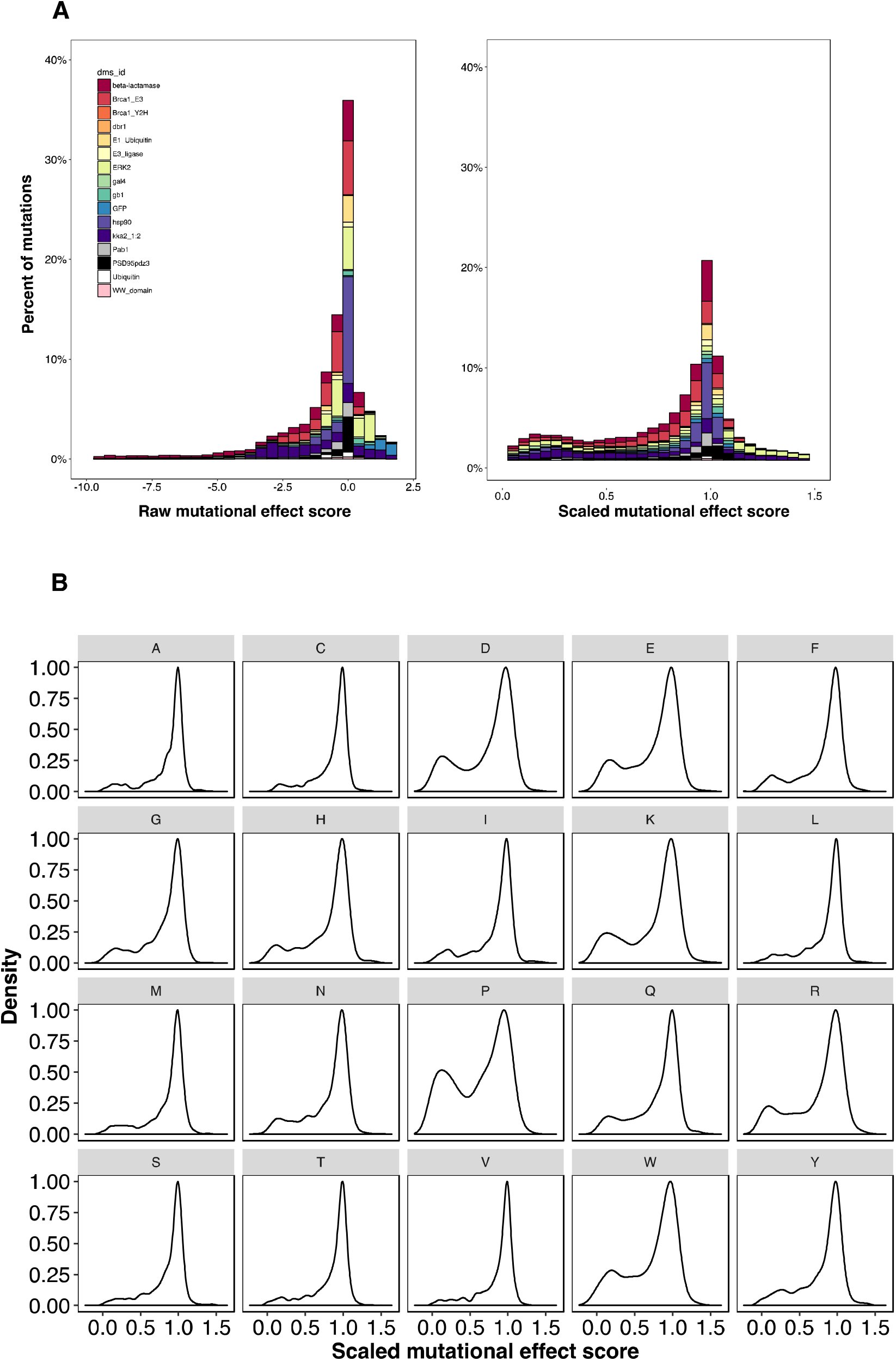
We curated large-scale mutagenesis data sets describing the effects of 34,373 mutations at 2,236 positions in fourteen proteins. To facilitate comparisons between each data set, we rescaled mutational effect scores for each protein by subtracting the median mutational effect score of all synonymous mutations in that protein from each nonsynonymous mutational effect score and then dividing that difference by the median of the bottom 1% of mutational effect scores (see **Methods**). **(A)** Stacked histograms of the original scores (**left panel**) and rescaled scores (**right panel**) are shown. **(B)** Density plots of the scaled mutational effect scores for each amino acid substitution are shown.

**Figure S2.**
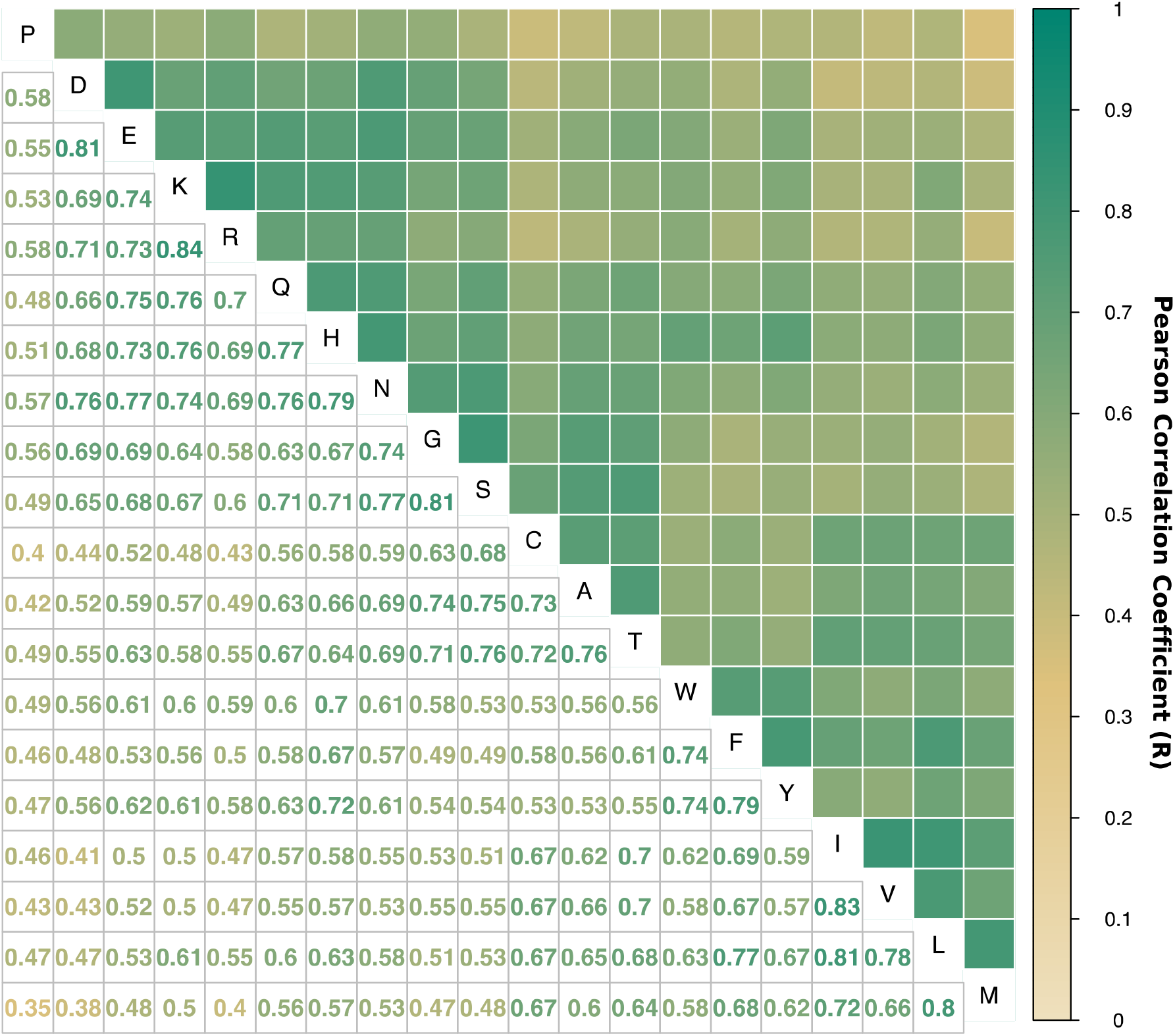
For each substitution, Pearson correlation coefficients were calculated for the mutational effect scores of that substitution with every other substitution at each position. A correlation plot of these Pearson coefficients is shown. Color indicates the Pearson correlation coefficient ranging from 0 (light brown) to 1 (green).

**Figure S3.**
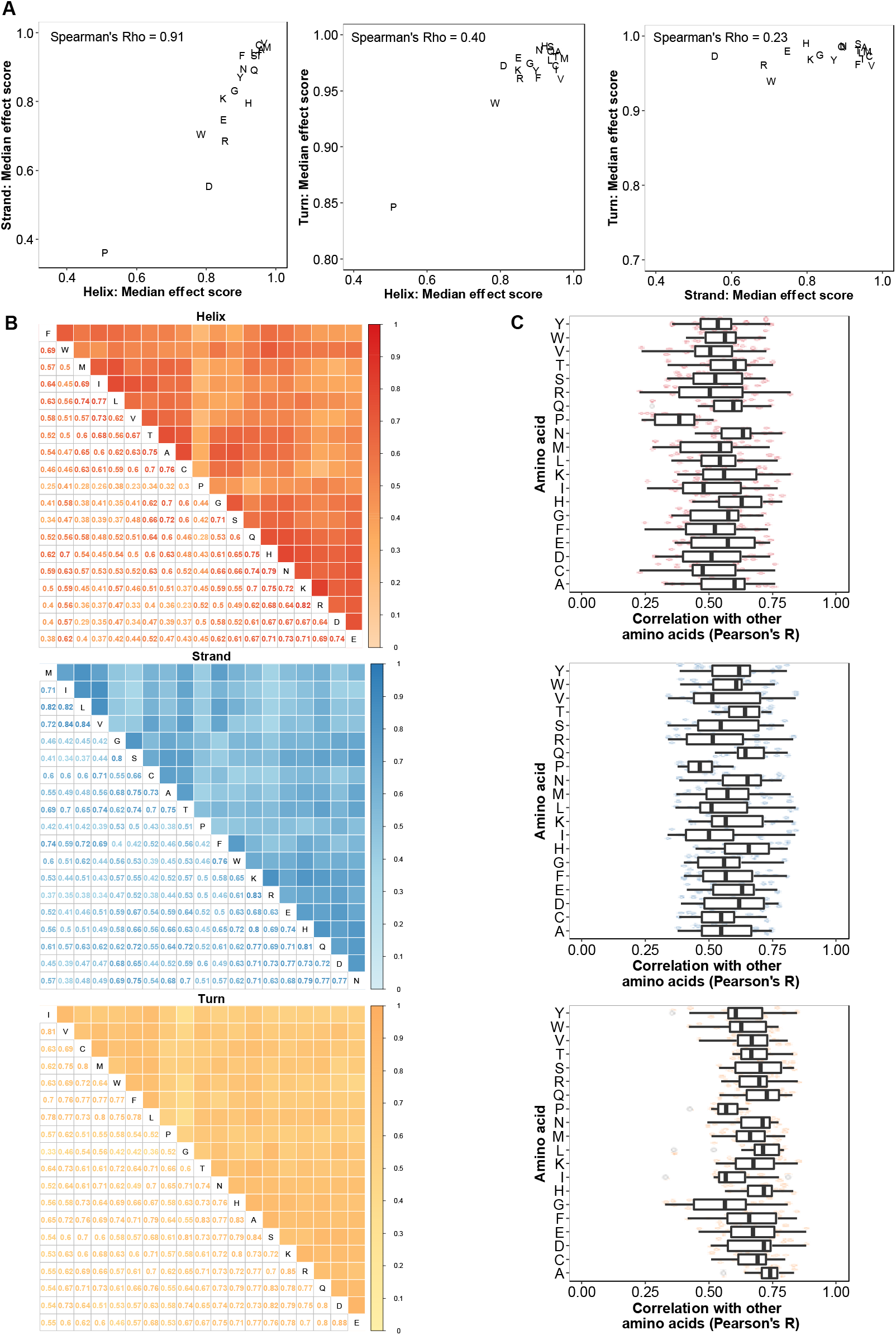
**(A)** For each amino acid substitution, the median mutational effect score was calculated. The correlation between the median mutational effects for each substitution in helices, strand and turns are shown in scatterplots, and Spearman’s Rho indicates the degree of rank correlation within each scatterplot. **(B)** Pearson correlation coefficients were calculated for the mutational effects of each substitution with every other substitution at every position. The Pearson correlation coefficient plots are shown separately for α-helices (top), β-sheets (middle), and turns (bottom). **(C)** Boxplots show the distribution of Pearson correlation coefficients for each amino acid type in three structural contexts.

**Figure S4.**
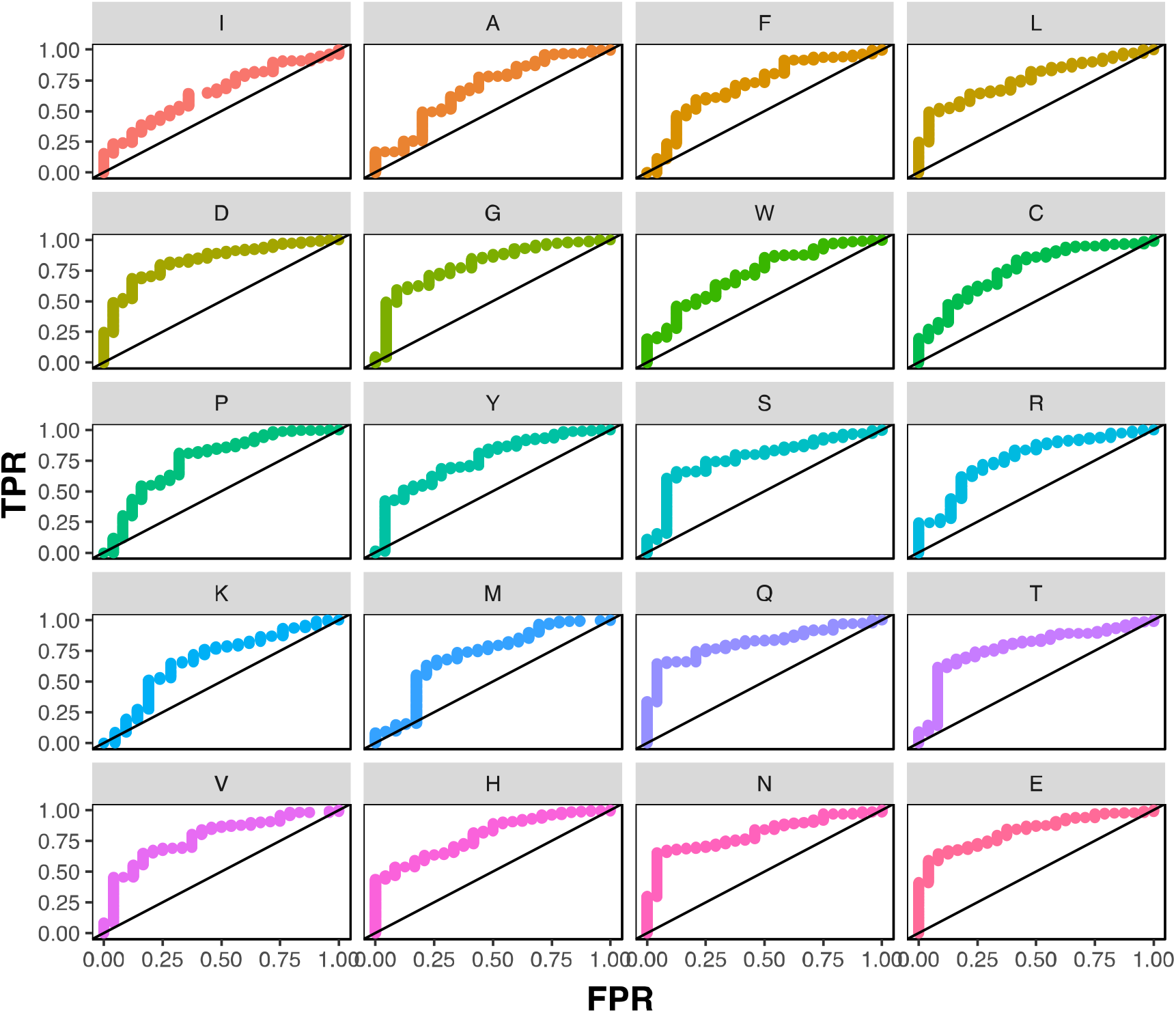
A mutational effect threshold was defined such that positions with a mutational effect score below the threshold were classified as “interface,” whereas positions with a mutational effect score above the threshold were classified as “non-interface.” ROC curves were generated by varying this threshold for each amino acid type in the four proteins with protein or DNA ligand-bound structures (hYAP65 WW domain, PSD95 pdz3 domain, Gal4 and BRCA1 RING domain (BARD1 binding)).

**Figure S5.**
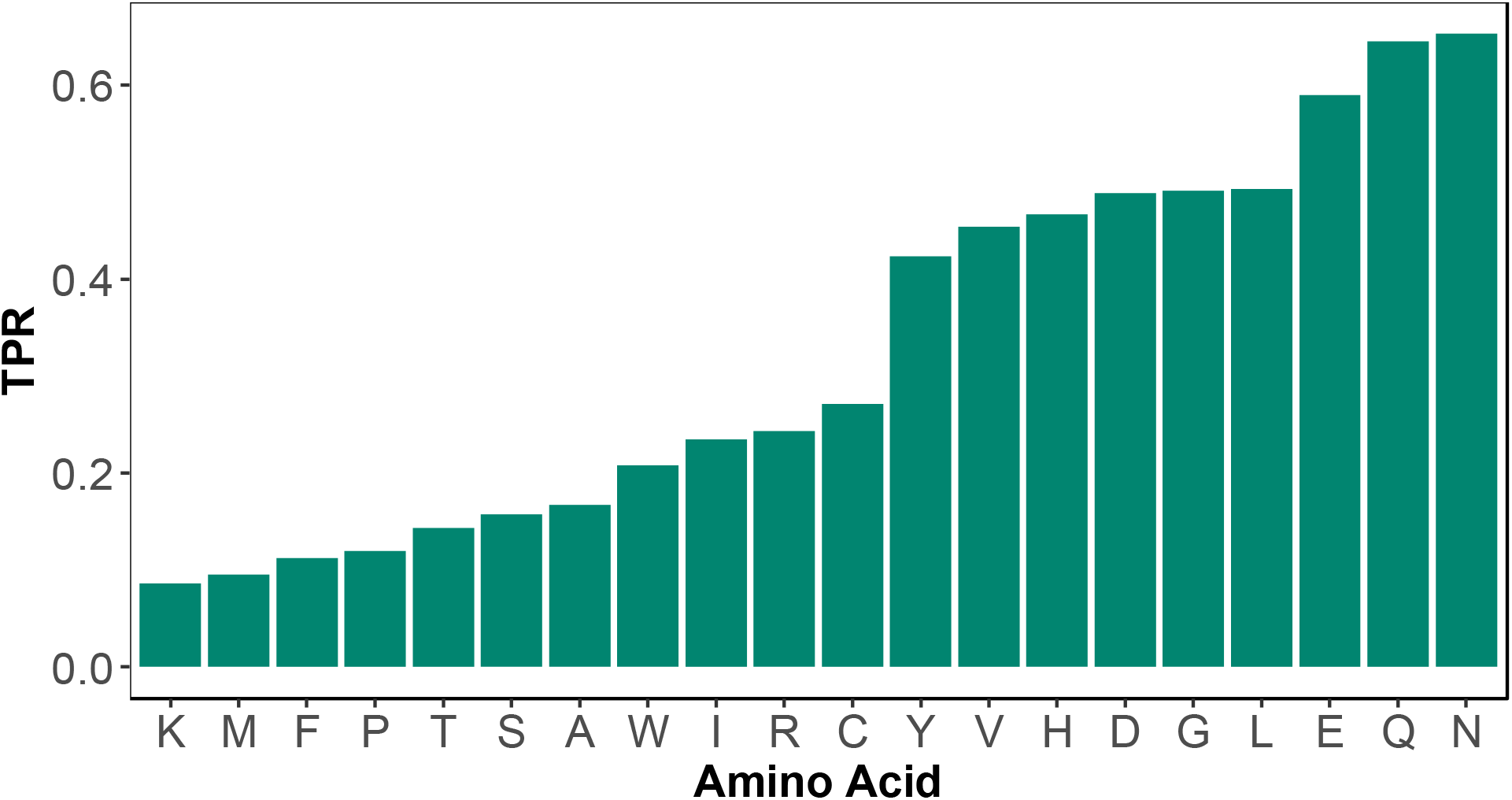
A mutational effect threshold was defined such that positions with a mutational effect score below the threshold were classified as “interface,” whereas positions with a mutational effect score above the threshold were classified as “noninterface.” A barplot shows each amino acid substitution’s true positive rate (TPR) for detecting interface positions at a fixed, 5% non-interface position false positive rate.

**Table S1.** A table showing sample size, *p*-value and Bonferroni corrected *p*-value for paired, two-sided Wilcoxon rank sum tests of the position median effect scores versus each amino acid substitution’s effect scores. This analysis was restricted to the 882 positions where the effects of all 19 possible substitutions were scored.

